# Immunotropic aspect of the *Bacillus coagulans* probiotic action

**DOI:** 10.1101/088757

**Authors:** T.V. Bomko, T. N. Nosalskaya, T. V. Kabluchko, Yu. V. Lisnyak, A. V. Martynov

## Abstract

Immunotropic aspect of the *Bacillus coagulans* probiotic action

**Objective:** Currently, probiotics are increasingly used as the alternative to antibiotics as well as the preventive measures in humans. In particular, probiotics occupy a key position in the treatment of antibiotics-associated intestinal dysbiosis. A spore-forming microorganism lactobacillus *Bacillus coagulans* is one of the most promising probiotics. However, some of its pharmacological effects remain poorly understood.

**Aim.:** This study is aimed at investigation of the effect of *Bacillus coagulans* (Laktovit Forte) on the intestinal dysbiosis syndrome in mice caused by streptomycin against the background of cyclophosphamide-induced cellular immunodeficiency.

**Methods.:** Pharmacological method: mouse model in vivo with immunodeficiency caused by cyclophosphamide.

**Key findings.:** In mice with colitis caused by streptomycin treatment, the administration of *Bacillus coagulans* (Laktovit Forte medicinal product) resulted in an antidiarrheal effect, normalization of gastrointestinal motility, and prevention of the animals’ weight loss. Given the cyclophosphamide-induced immunosuppression and streptomycin-associated diarrhoea, the immunity was completely restored only under the action of *Bacillus coagulans*.

**Conclusions.:** According to all parameters, *Bacillus coagulans* has been proved to be more effective as compared to the Linex Forte reference product containing lacto‐ and bifidobacteria.

## Introduction

Various factors of external and internal environment can significantly affect the composition of intestinal microflora that may disturb the normal course of physiological processes and even result in severe pathological states, one of which is dysbiosis syndrome. The most common cause of intestinal dysbiosis syndrome is the use of antibiotics, which directly suppress the intestinal microorganisms’ vital activity and significantly change a “microbial landscape” of the gastrointestinal tract ^[1,2]^.

Disorders of the fermentation of carbohydrates, proteins and lipids, caused by the dysbiotic microflora, result in the strengthening of zymosis and putrefaction processes in different parts of the digestive tract that, in turn, stimulates an increasing toxic substances formation, including Gram-negative bacteria’s endotoxins, deterioration of the normal flora’s main physiological functions, and growth of the opportunistic pathogens’ population. As a result, there is an increase of the structural and functional changes in the digestive organs: the inflammatory process intensification, the membrane digestion disorder and intestinal malabsorption, and dysbiosis syndrome development ^[3]^.

The mechanism of antibiotic-associated diarrhoea is considered to be related to the growth of the pathogenic microorganisms producing toxin A. The toxin A induces COG-2 expression in colonocytes, that is the development of inflammatory response and diarrhoea, activates nuclear factor NF-κB, assisting in the chemokines production and taking part in the colonocytes apoptosis, and assists in the production of activator protein-1 (AP-1), involved in the interleukin IL-8 production ^[4]^.

Currently, probiotics occupy a key position in the treatment of antibiotics-associated intestinal dysbiosis. To be included in the group of probiotics the microorganisms should meet the following criteria: 1) to survive the passage through the gastrointestinal tract, i.e. to be acid and bile resistant; 2) to adhere to intestinal epithelial cells; 3) to grow fast, colonize the intestinal tract, to persist there, and then to leave the body; 4) to stabilize the intestinal microflora; 5) to have no signs of pathogenicity; 6) to maintain viability both in food and during the pharmacopoeial drugs manufacturing ^[5,6]^. Some representatives of spore-forming bacteria of the genus *Bacillus* satisfy these requirements to a greatest extent. These microorganisms are capable to prevent intestinal disorders, acting in some cases more effectively than the conventional probiotics containing bifidobacteria and lactobacilli ^[7,8]^. Lactobacilli *Bacillus coagulans*, a spore-forming microorganism, is one of the most promising candidate for probiotics. When in the form of spores, it is resistant to the manufacturing processes and storage, and is not lysed by the gastric fluid and bile. In the duodenum, the spores of *Bacillus coagulans* can germinate into vegetative bacteria showing probiotic activity; then they are excreted from the body without alteration of the individual human microflora’s composition ^[9]^. *Bacillus coagulans* is not merely a component for dietary supplements. In recent years, a number of drugs containing this bacterium have entered the global pharmaceutical market and have already proven their clinical efficacy ^[10]^.

However, some of the pharmacological effects of *Bacillus coagulans* remain poorly understood. In particular, it concerns its strong immunomodulatory effect, antiviral activity, and stimulation of the normal intestinal microflora’s rapid recovery ^[11]^. Experimental and clinical studies have confirmed the immunomodulatory effect of the *Bacillus coagulans*. For instance, when the blood cells in healthy human volunteers were exposed to adenovirus and influenza A, the administration of probiotic significantly increased the production of the markers of CD3-CD69+cells, IL-6 and IL-8 interleukins, *γ*-interferon, and TNF-α^[12]^.

*Bacillus coagulans* showed its efficacy against neonatal rotavirus-induced diarrhoea ^[13]^; it also significantly weakened the rheumatoid arthritis’ symptoms intensity ^[14]^. These effects can be related to a direct immunomodulatory action of *Bacillus coagulans* spore forms ^[15]^. During the *in vitro* experiments the metabolites and cell walls of *Bacillus coagulans* intensified the phagocytosis (the polymorphonuclear leukocytes’ population increased, including the phagocytic cells), the migration of polymorphonuclear leukocytes (spontaneous migration and the migration caused by the bacterial peptide f-MLP), and the inhibited migration induced by pro-inflammatory interleukin-8 and leukotriene LTB-4. Both components of *Bacillus coagulan*s induced the expression of killer cells (CD69 marker) and affected the production of cytokines: inhibited IL-2 production, enhanced production of IL-4, IL-10, and, especially, IL-6. These effects were aimed at the stimulation of the B lymphocytes’ proliferation and differentiation, and at the anti-inflammatory action. In addition, the probiotic significantly increased the *γ*-interferon production ^[16]^. The experiments showed the similar species *Bacillus subtilis* spores’ ability to penetrate into the Peyer’s patches and interact with the gut-associated lymphoid system, and to accumulate in macrophages and germinate therein ^[17]^. Although the spores’ germination within macrophages is not terminated by the formation of vegetative bacteria, the immune system interprets such spores’ behaviour as an invasion ^[14]^.

The *Bacillus coagulans* bacterium is called “the king of probiotics” due to its high stability in the gastrointestinal tract, non-toxic action, as well as high but not yet fully understood pharmacological activity.

This study aims to investigate the effect of *Bacillus coagulans* (Laktovit Forte medicinal product) on the clinical manifestations of the intestinal dysbiosis syndrome in mice caused by the administration of streptomycin against the background of cyclophosphamide-induced cellular immunodeficiency.

## Materials and methods

### Animals, diarrhoea, immunodeficiency

Animal studies were performed according to the European Convention for the Protection of Vertebrate Animals used for Experimental and other Scientific Purposes ^[18,19]^. Bioethical aspects of the studies have been approved by the Committee on Bioethics of the I. Mechnikov Institute of Microbiology and Immunology of the National Academy of Medical Sciences of Ukraine State Institution in accordance with the international standards ^[20,21]^.

Non-inbred male mice weighing 24 to 25 g, were kept in standard laboratory conditions (20– 22 °C, 14 h/10 h light/dark cycle, 65% humidity) supplied with food and water ad libitum. The standard mouse food pellets contained a balanced diet. The procedures followed were in accordance with the institutional guidelines.

To simulate the dysbiosis syndrome the mice were intragastrically administered streptomycin (STM) (Arterium, UA, serial No. 125894) at a dose 2 g/kg of body weight for 9 days ^[22,23]^.

Immune status disorders were simulated by a single subcutaneous injection of cyclophosphamide (CFA) (Endoxan, Baxter oncology GmbH, Halle/Germany, Serial No. 4J035F) at a dose of 250 mg/kg. The CFA subcutaneous injection ensures the development of a long-term (7-10 days) immunocompromised state in mice ^[24]^. The mice were administered CFA 1 h before the streptomycin <u>single-dose administration</u>.

Laktovit Forte (LAF) (“Mili Healthcare”, Great Britain, Serial No. LF171) contained 120 × 10^6^ spores of *Bacillus coagulans*, 0.0015 g folic acid, and 15 μg cyanocobalamine. Animals were intragastrically administered LAF at a dose 46 mg/kg. Before administering LAF, the capsule content was suspended in distilled water. To remove vitamins, the suspension was dissolved and centrifuged for 25 minutes at <u>5 * 10^3^ g (r = 7 cm, rpm = 9 * 10^3^)</u>. Supernatant containing vitamins was discarded and the precipitate was re-suspended and administered to animals.

Used as a control drug was Linex Forte (LIF) (“Sandoz”, EU, Serial No. FK5074) containing only non-spore-forming bacteria: 1 × 10^9^ CFU of *Lactobacillus acidophilus* (LA-5), 1 × 10^9^ CFU of *Bifidobacterium animalis subsp. Lactis BB-12*). LIF was administred *per os* at a dose of 42 mg/kg. Both drugs were administered for 7 days after a 5-day course of antibiotic. The above experimental protocol was based on our previous studies, which showed that in case of streptomycin-induced pathology the faeces liquefaction (diarrhoea) was observed since the 6^th^ day of the drug administration.

Animals were randomized into 7 groups, each comprising 12 mice:

1. – control (intact);
2. – pathology control (STM);
3. – pathology with immunosuppression (STM and CFA);
4. – STM + LAF;
5. – pathology with immunosuppression (CFA) + LAF treatment;
6. – STM + LIF;
7. – pathology with immunosuppression (CFA) + LIF treatment.

Animals, weighed prior to and on the 12^th^ day of the experiment, were kept in cages <u>in the individual compartments</u>. Monitored during the experiment were the characteristics of faeces. To assess the characteristics (consistency) of faeces used was a five-point scale, where: 1 – soft, 2 – semi-liquid, 3 – liquid without mucus, 4 – liquid with mucus, and 5 – mucus.

To estimate an antidiarrheal effect (ADE), the percentage of animals without diarrhoea was calculated by the following equation:

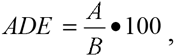

where A is a number of animals without diarrhoea; B is a number of animals in the group.

To determine the gastrointestinal motility, the animals were given a charcoal “label” (0.5 ml of 10% activated charcoal suspension per mouse) at the end of the experiment (on the 12^th^ day). Forty minutes later the animals were sacrificed by cervical dislocation, with their intestine and spleen being isolated. To determine the cytokines level and the number of macrophages and lymphocytes, the blood was collected from the animals’ hearts.

The activity of intestinal peristalsis was evaluated by the share of the intestine filled with the charcoal “label”. The normalization of intestinal motor activity (NIMA) was calculated in percent by the equation:

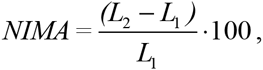

where L_1_ and L_2_ are the lengths of the intestine filled with “label” in the control group and in the experimental group respectively.

## Immunological methods

### NO and cytokine production assay.

Isolated were murine peritoneal macrophages and splenic T-lymphocytes in order to measure the TNF production using a sandwich enzyme immunoassay (ELISA) technique with the ELISA Kit (Mouse TNF-a ELISA Kit, PharMingen, BD Biosciences, San Jose, CA, USA), performed according to ^[25]^.

Macrophages, 10^6^ cells/ml, were cultivated in 24-well stripped plates (BioRad, Hannover, EU) in Medium 199 with a methyl red pH-indicator at 37 °C in humidified atmosphere containing 5% CO_2_. The medium was supplemented with 25mM HEPES, 3% foetal bovine serum. Supernatants were collected at 21 h and assayed for NO_2_. The Griess reagent (containing 1% sulphanilamide/0.1% naphthylethylene diamine dihydrochloride and 2.5% H_3_PO_4_ (1 : 1)) was used to estimate NO2, which served as indicator of NO production. After cultivating, a 100 ml of each sample was placed into a 96-well plate with a 100 ml of the Griess reagent. Ten minutes later, the optical density was measured on the trip reader (Stat Fax 303 plus, Awareness technology Inc., USA) at 590 nm ^[26]^. <u>The same reader was used for all ELISA studies</u>.

### IL-2 assay

Spleen lymphocytes, 1.5 × 10^6^ cells/ml, were cultivated for 72 h in 24-well stripe plates in 199 medium at 37 °C in humidified atmosphere containing 5% CO_2_. The medium was supplemented with 25 mM HEPES, 5% foetal bovine serum, and 5 mg/ml phytohemagglutinin (FHA). The IL-2 concentration in the supernatant of the PHA-stimulated cells was measured by the ELISA using rabbit anti-murine IL-2 polyclonal antibody (BioRad, AbD Serotec, Raleigh, NC, USA) and goat anti-rabbit IgG Biotin Conjugate (Agrisera Antibodies, Vännäs, SWEDEN). After staining with ABTS, the optical density was measured on the strip reader at 405 nm ^[27]^.

### Statistical analysis

All the experimental data were processed by the method of variation statistics. Calculation of statistical significance in the case of nominal variables was performed using ANOVA and ANOVA for experiments with the repeated measures. Statistical significance was calculated by Student’s t-test in Bonferroni modification ^[28]^.

## Results and discussion

### The antidiarrheal effect of Laktovit Forte and Linex Forte

On the 6^th^ day of streptomycin administration, the faeces softening was observed in all animal groups (Table 1). The most pronounced diarrhoea was registered in the pathology control groups (the 2 and 3) from the 9^th^ to 12^th^ day of the experiment.

**Table 1.** Development of diarrhoea in mice treated with streptomycin, Laktovit Forte and Linex

| Mice group (12 mice each) | The faeces consistency on the observation day (from 5th to 12th) <sup>a</sup> |  |  |  |  |  |  |  |
| --- | --- | --- | --- | --- | --- | --- | --- | --- |
|  | 5 | 6 | 7 | 8 | 9 | 10 | 11 | 12 |
| 1 | 1.0 ± 0.00 | 1.0 ± 0.00 | 1.0 ± 0.00 | 1.0 ± 0.00 | 1.0 ± 0.00 | 1.0 ± 0.00 | 1.0 ± 0.00 | 1.0 ± 0.00 |
| 2 | 1.0 ± 0.00 | 1.92 ± 0.23 | 2.50 ± 0.15 | 3.67 ± 0.22 | 4.67 ± 0.14 | 4.75 ± 0.13 | 4.75 ± 0.13 | 4.67 ± 0.14 |
| 3 | 1.0 ± 0.00 | 1.77 ± 0.19 | 3.00 ± 0.21 | 4.25 ± 0.18 | 4.83 ± 0.11 | 4.83 ± 0.11 | 4.75 ± 0.15 | 4.58 ± 0.15 |
| 4 | 1.0 ± 0.00 | 1.58 ± 0.13 | 2.20 ± 0.11 | 2.17 ± 0.24 | 2.00 ± 0.21 | 1.67 ± 0.19 | 1.42 ± 0.15 <sup>b</sup> | 1.25 ± 0.13 <sup>b</sup> |
| 5 | 1.0 ± 0.00 | 2.00 ± 0.17 | 2.42 ± 0.19 | 2.08 ± 0.23 | 2.00 ± 0.21 | 2.0 ± 0.174 | 1.750 ± 0.218 | 1.58 ± 0.19 <sup>b</sup> |
| 6 | 1.0 ± 0.00 | 1.75 ± 0.18 | 2.67 ± 0.22 | 2.67 ± 0.28 | 2.75 ± 0.33 | 2.667 ± 0.284 | 2.583 ± 0.288 | 2.08 ± 0.31 |
| 7 | 1.0 ± 0.00 | 2.0 ± 0.25 | 3.0 ± 0.302 | 3.17 ± 0.34 | 3.17 ± 0.39 | 3.17 ± 0.32 | 2.75 ± 0.30 | 2.50 ± 0.42 |

<sup>a</sup> – The differences of the faeces consistency in a group on a day of observation, relative to the 5<sup>th</sup> day, were statistically significant (p≤0.05).
<sup>b</sup> – The differences between experimental groups and control (group 1) on a day of observation were statistically significant (p≤0.05).

During this period, the diarrhoea was manifested as a shapeless liquid stool with mucus and often as a mucus only, which corresponded to 4-5 in the five-point scale. No significant difference was observed between the animal groups, treated with streptomycin only (the group 2) and treated with streptomycin plus cyclophosphamide (the group 3).

When LAF was administered to the animals (the group 4), their diarrhoea weakened and was mildly manifested on the 9^th^ day: their faeces were semi-shapeless and, in some cases, semi-liquid. However, in the animal group 5, comprising the mice that were administered with LAF against the background of immunosuppression, the diarrhoea was significantly inhibited (p≤0.05).

In mice treated with LIF (the groups 6 and 7) the diarrhoea was more prominent than in the groups 3 and 4, but less prominent than in the group 2 (pathology control).

The parameters of the antidiarrheal effect (table 2) also revealed a high activity of LAF: ADE values amounted to 75%. In the group of mice with immune imbalance, the probiotic’s antidiarrheal effect was less, ADE values amounted to 50%, indicating that the antidiarrheal action of *Bacillus coagulans* (LAF) possibly depends on the immune system.

**Table 2.** The antidiarrheal effect of Laktovit Forte and Linex Forte

| Mice group | ADE, % |
| --- | --- |
| 1 (Intact – first control group) | - |
| 2 (STM – second control group) | - |
| 3 (STM+CFA – third control group ) | - |
| 4 (STM +LAF – first training group) | 75.0 % |
| 5 (STM +CFA+LAF – second training group) | 50.0% |
| 6 (STM +LIF – third training group) | 41.7% |
| 7 (STM +CFA+LIF – fourth training group) | 33.3% |

The antidiarrheal effect of LIF was 41.7% in the group treated with streptomycin, and 33.3% in the group with diarrhoea and immunosuppression, which also may indicate a certain immunomodulatory role of bacteria from the LIF. The effect of probiotics on the mice gastrointestinal motility showed similar tendencies (Table 3).

**Table 3.** Effect of probiotics on the gastrointestinal motility of mice with streptomycin-induced

| Mice group | Intestine length (cm): |  | % of the total intestine length | Normalization of gastrointestinal motility,% |
| --- | --- | --- | --- | --- |
|  | total | filled with “label” |  |  |
| 1 | 39.17 ± 1.60 | 11.22 ± 0.31 | 28.7 | - |
| 2 | 39.08 ± 1.62 | 24.92 ± 0.86 | 63.8 | - |
| 3 | 38.25 ± 1.17 | 25.22 ± 0.62 | 65.9 | - |
| 4 | 37.73 ± 0.87 | 14.67 ± 0.58 <sup>a,b</sup> | 38.8 | 61.1 |
| 5 | 39.01 ± 0.95 | 17.96 ± 0.68 <sup>a,b</sup> | 46.0 | 54.0 |
| 6 | 39.34 ± 1.06 | 21.85 ± 0.54 <sup>a</sup> | 55.5 | 44.5 |
| 7 | 41.40 ± 0.98 | 22.37 ± 0.85 | 54.0 | 45.9 |

<sup>a</sup> - Statistically significant difference compared to the group 2 and 3 (p<0.05)
<sup>b</sup> - Statistically significant difference compared to the reference drug (p<0.05)

The streptomycin administration during 9 days had increased the intestinal peristalsis, which was evidenced by a significant increase of the “labelled” intestine share: up to 63.8% against 28.7% in the intact control group. This testifies to the fact that the streptomycin administration enhanced the gastrointestinal motility by 2.4 times. The enhanced intestine peristalsis was accompanied by a more rapid evacuation of the bowel content, namely by the increased number of defecations and by the faeces softening. A similarly enhanced intestine peristalsis was observed in the animal group, treated with STM and CFA.

The administration of LAF against the background of pathology resulted in the decreased intestine peristalsis activity, with its normalization amounting to 61%. In the mice group 5, the normalization of the intestine peristalsis activity under the influence of probiotic was slightly lower and amounted to 54%.

The effect of LIF was significantly less prominent: the intestine peristalsis activity normalization in the mice groups 6 and 7 was only 44.5% and 45.9% respectively.

The progression of diarrhoea had a negative effect on the animals’ weight (Table 4). In the mice group 1 (intact) the weight increased by 10%, while in the groups 2 and 3 (pathology and one with the immunosuppression) there was observed a decrease in the weight up to 11.5% and 12.8% respectively.

**Table 4.** Effect of the streptomycin-induced diarrhoea and probiotics on the mice weight

| Mice group | Body weight (g) |  |
| --- | --- | --- |
|  | Before experiment | On the 12<sup>th</sup> day |
| 1 | 20.83 ± 0.48 | 22.917 ± 0.319 * |
| 2 | 21.00 ± 0.25 | 18.58 ± 0.41 * |
| 3 | 20.83 ± 0.41 | 18.17 ± 0.63 * |
| 4 | 20.75 ± 0.48 | 22.83 ± 0.71 * |
| 5 | 20.75 ± 0.64 | 21.30 ± 0.48 |
| 6 | 20.75 ± 0.41 | 21.75 ± 0.41 * |
| 7 | 20.75 ± 0.96 | 21.58 ± 0.25 |

\* - Statistically significant difference compared to the weight before the experiment ( $p \leq 0.05$ )

The administration of streptomycin in high doses in mice is known to cause the inflammatory bowel disease (colitis) and to decrease their resistance to infection ^[22]^. The progression of diarrhoea, the parietal digestion disorder, and absorption disorder result in the weight loss. The administration of LAF against the background of pathology assisted in the parameters improvement and, accordingly, in gaining the weight.

### The effect of LIF and LAF on the mice immunity

The effect of streptomycin on the mice immune system was not prominent (Table 5), and was accompanied by a significant reduction in the IL-2 production.

**Table 5.**
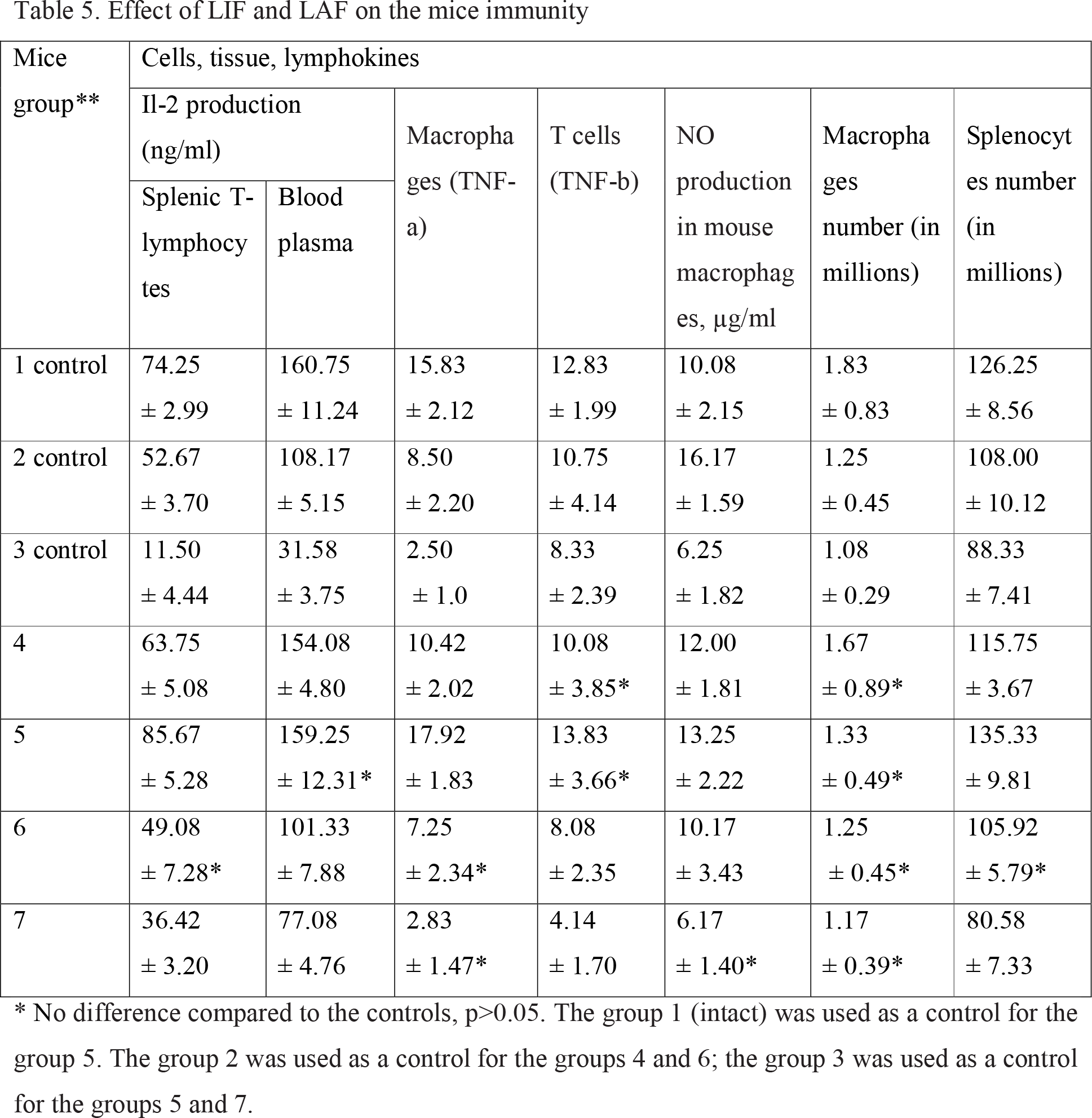
Effect of LIF and LAF on the mice immunity

| Mice group** | Cells, tissue, lymphokines |  |  |  |  |  |  |
| --- | --- | --- | --- | --- | --- | --- | --- |
|  | Il-2 production (ng/ml) |  | Macrophages (TNF-a) | T cells (TNF-b) | NO production in mouse macrophages, µg/ml | Macrophages number (in millions) | Splenocytes number (in millions) |
|  | Splenic T-lymphocytes | Blood plasma |  |  |  |  |  |
| 1 control | 74.25<br>± 2.99 | 160.75<br>± 11.24 | 15.83<br>± 2.12 | 12.83<br>± 1.99 | 10.08<br>± 2.15 | 1.83<br>± 0.83 | 126.25<br>± 8.56 |
| 2 control | 52.67<br>± 3.70 | 108.17<br>± 5.15 | 8.50<br>± 2.20 | 10.75<br>± 4.14 | 16.17<br>± 1.59 | 1.25<br>± 0.45 | 108.00<br>± 10.12 |
| 3 control | 11.50<br>± 4.44 | 31.58<br>± 3.75 | 2.50<br>± 1.0 | 8.33<br>± 2.39 | 6.25<br>± 1.82 | 1.08<br>± 0.29 | 88.33<br>± 7.41 |
| 4 | 63.75<br>± 5.08 | 154.08<br>± 4.80 | 10.42<br>± 2.02 | 10.08<br>± 3.85* | 12.00<br>± 1.81 | 1.67<br>± 0.89* | 115.75<br>± 3.67 |
| 5 | 85.67<br>± 5.28 | 159.25<br>± 12.31* | 17.92<br>± 1.83 | 13.83<br>± 3.66* | 13.25<br>± 2.22 | 1.33<br>± 0.49* | 135.33<br>± 9.81 |
| 6 | 49.08<br>± 7.28* | 101.33<br>± 7.88 | 7.25<br>± 2.34* | 8.08<br>± 2.35 | 10.17<br>± 3.43 | 1.25<br>± 0.45* | 105.92<br>± 5.79* |
| 7 | 36.42<br>± 3.20 | 77.08<br>± 4.76 | 2.83<br>± 1.47* | 4.14<br>± 1.70 | 6.17<br>± 1.40* | 1.17<br>± 0.39* | 80.58<br>± 7.33 |

\* No difference compared to the controls, $p > 0.05$ . The group 1 (intact) was used as a control for the group 5. The group 2 was used as a control for the groups 4 and 6; the group 3 was used as a control for the groups 5 and 7.

The macrophages and T-cells also significantly decreased the TNF production, and at the same time the macrophages increased the NO production. The number of splenic lymphocytes and macrophages significantly decreased, but remained not lower than the life-threatening level for the animals. The combination of CFA and STM significantly influenced on both the exacerbation of intestinal dysbiosis and the immune system (the group 3): the T-lymphocytes’ production of the IL-2 in spleen and in blood plasma decreased more than five-fold. The TNF-a level had similarly decreased. Also, the level of TNF-b, though significantly decreased, was not so essential as in the group 1. The NO, produced by macrophages (6,25 ± 1,82 μg/ml against 10,08 ± 2,15 μg/ml), decreased almost two-fold in the group 3, comprising animals that were administered with cyclophosphamide and streptomycin. Among the experimental groups, the best results of the immune recovery to nearly normal levels were observed in the group 5, where LAF was administered against the background of CFA. The level of splenic T-lymphocytes was even higher in the control group (although not statistically significant, p<0.1). The parameters in the animal groups without the artificial immunosuppression (the groups 4 and 6) were close to those of the control group 2 (streptomycin). But it was only the parameters from the group 4 (LAF introduction on the background of STM) that differed significantly (p<0.05) and, according to all parameters, were similar to the ones from the control group 2 (streptomycin). The only inexplicable phenomenon, going beyond the immunosuppression concept, is the IL-2, produced by the splenic lymphocytes, exceeding the control values in the group 5 (with induced immunosuppression). We attribute this fact to the non-uniform effect of CFA on different parts of the immune system and their non-synchronous stimulation by the *B. coagulans* spore form. In contrast to the LAF, the LIF had stimulated the immunity compared to the control group 3, but could not recover it even up to the level of the control group 2 (with streptomycin dysbiosis).

## Conclusions

In mice with colitis, caused by streptomycin treatment, the administration of *Bacillus coagulans* (Laktovit Forte medicinal product) resulted in an antidiarrheal effect (diarrhoea syndrome decreased by 75%), normalization of gastrointestinal motility, and prevention of the animals’ weight loss. The less pronounced antidiarrheal effect was demonstrated in case of cyclophosphamide-induced immunosuppression, which suggests that the role of immune system is important in the probiotic’s mechanism of action. The *Bacillus coagulans* had normalized both the quantitative parameters of the immune system (the number of splenic lymphoocytes, macrophages, and T-lymphocytes) and the cells’ functional activity (production of IL-2 and TNF). Given the cyclophosphamide-induced immunosuppression and streptomycin-associated diarrhoea, the immunity recovers completely only under the effect of *Bacillus coagulans*.

Thus, by all parameters, *Bacillus coagulans* was shown to be more effective compared to the Linex Forte reference product containing lacto‐ and bifidobacteria.

## Declarations

Conflict of interest

Authors declare that they have no conflict of interests to disclose.

## Funding

We gratefully acknowledge the National Academy of Medical Sciences of Ukraine (NAMSU, Kiev, Ukraine), for the award of research fellowships. Financial support from NAMS, NAMSU131/2016 0116U000866, Government of Ukraine, is acknowledged. In addition, we gratefully acknowledge “Mili Healthcare Ltd.” for the technical support and consultation, and for the samples of probiotics granted.

